# Data-Driven Modeling of Resource Distribution in Honeybee Swarms

**DOI:** 10.1101/2020.05.13.090704

**Authors:** Golnar Gharooni Fard, Elizabeth Bradley, Orit Peleg

## Abstract

Trophallaxis is the mutual exchange and direct transfer of liquid food among eusocial insects such as ants, termites, wasps, and bees. This process allows efficient dissemination of nutrients and is crucial for the colony’s survival. In this paper, we present a data-driven agent-based model and use it to explore how the interactions of individual bees, following simple, local rules, affect the global food distribution. We design the rules in our model using laboratory experiments on honeybees. We validate its results via comparisons with the movement patterns in real bees. Using this model, we demonstrate that the efficiency of food distribution is affected by the density of the individuals, as well as the rules that govern their behavior: e.g., how they move and whether or not they aggregate. Specifically, food is distributed more efficiently when donor bees do not always feed their immediate neighbors, but instead prioritize longer motions, sharing their food with more-distant bees. This non-local pattern of food exchange enhances the overall probability that all of the bees, regardless of their position in the colony, will be fed efficiently. We also find that short-range attraction improves the efficiency of the food distribution in the simulations. Importantly, this model makes *testable* predictions about the effects of different bee densities, which can be validated in experiments. These findings can potentially contribute to the design of local rules for resource sharing in swarm robotic systems.

## Introduction

Autonomous, adaptive and robust interaction networks allow super-organisms to exchange information and resources efficiently and effectively between their individual building blocks (Sumpter, 2010). These networks play critical roles in the function of these entities. In honeybees (Seeley, 2009), for instance, communication networks allow sensing and reproduction of tactile signals, thus serving as “signal amplifiers” for information about the location of food sources. And bees do not just share information about the location of food; they also share food itself via regurgitation, essentially “charging” nestmates who do not have access to that resource (Robinson, 1992). This process, termed Trophallaxis LeBoeuf (2017) (shown in Fig. 2(c)), allows dissemination of nutrients through the colony and is crucial for its survival (Farina, 1996). There are subtleties in the trophallaxis process, though. Among other things, a bee that just finished feeding a nestmate faces a dilemma that is depicted in Fig. 1(a): should it feed another nearby nestmate or move to a new spot to feed a more-distant nestmate? To answer that question, and to identify an efficient set of local rules of behavior that guide food distribution, we construct an Agent-Based Model (ABM) of interactions via trophallaxis among autonomous agents. We conduct laboratory experiments to design and validate this model, then use it to explore different scenarios: different movement geometries, for instance, and agent densities.

**Figure 1:**
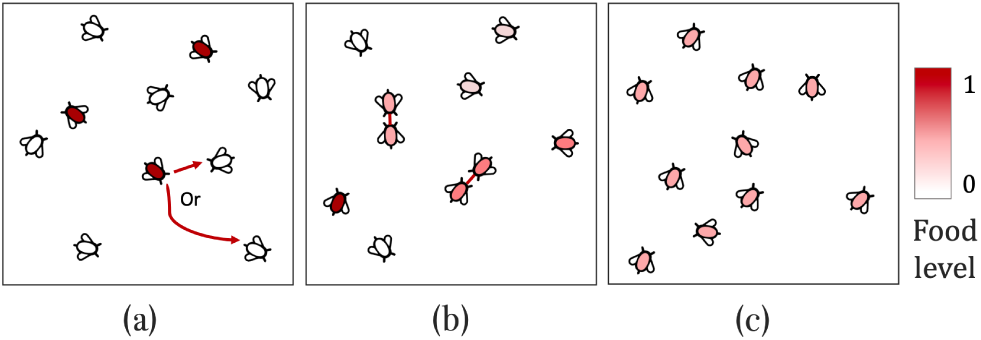
Agent-based model of trophallaxis behavior in honeybees. (a) At the beginning of the simulations, the fed (dark red) and the deprived (white) bees are scattered randomly in the simulation arena (b) Intermediate state of the system, showing two pairs of agents performing trophallaxis exchanges. (c) When food is distributed evenly across all agents, the model stops.

**Figure 2:**
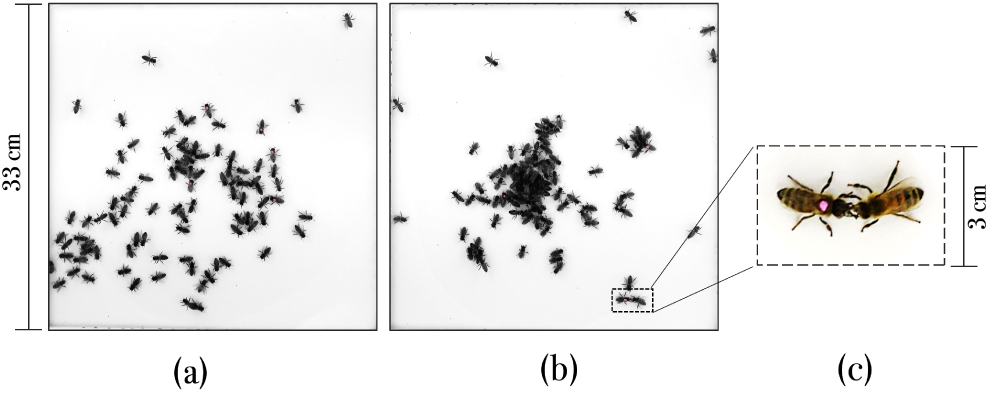
Experimental setup. Fed bees are marked with pink dots; unmarked bees are deprived of food until the beginning of the experiment. (a) A top view of the arena a few seconds after the introduction of the full bees; (b) View of the same experiment after 2 minutes; (c) An instance of trophallaxis.

Computational models of collective behavior provide quantitative and empirically verifiable accounts of how individual decisions and interactions can lead to the emergence of group-level outcomes (Goldstone and Janssen, 2005). In particular, agent-based models have been widely used in computational modeling of spatiotemporal dynamics, such as disease spread (Christakos et al., 2017), urban sprawl (Batty, 2013), traffic flows (Al-Dohuki et al., 2016), animal behavior (DeAngelis and Diaz, 2019), etc. As for modeling collective trophallactic systems, previous theoretical and computational approaches made significant progress by using tools from network theory to describe the network of trophallactic interactions in the colony (Gräwer et al., 2017), as well as tools from statistical physics to describe the food intake in a colony at the limits of infinite colony size and long time scales (Gräwer et al., 2017; Greenwald et al., 2018). These approaches provide valuable insight into the food-dissemination process but neglect the movement ecology of individual insects in physical space. Here, we attempt to fill this gap by using results from laboratory experiments to extend these models, making them a better match for natural behavior in honeybees.

Our work focuses specifically on the patterns of food distribution across the colony, tracking it over time as the model runs under different parameter conditions. Assuming a Correlated Random Walk (CRW) for the movement of the bee-agents in our model, as described in the following section, we look at the effect of several attributes of the simulations— density of agents and the width of turning angle distribution of the random-walk performed by individuals at each time step upon the efficiency of the food distribution across the group. We then perform a series of experiments with honeybees *Apis Mellifera* L., carefully monitoring and recording their positions, movements, and food distribution encounters. These experiments are described in the third section of the paper (“laboratory Experiments”). We use the empirical knowledge obtained from these experiments to design a data-driven model that is a good match for the experiments: serving as a reference to find physically realistic values for the free parameters in the model, for instance, and as a basis for validation of the simulation results. These results, as described in the penultimate section of this paper, enable us to finalize the design of the model, selecting appropriate values for a particularly influential parameter: the attraction range. Small values of this parameter—i.e., short-range attraction between bee-agents—increase the efficiency of food distribution. Simulations of this model suggest that a more tightly directed random walk increases the spatiotemporal efficiency of food distribution. We also find that the density of the bees in the simulation arena is highly influential in the results. The results of this study could potentially be used in designing a local rule set for efficient, self-organizing resource-distribution systems.

## Model Description

ABMs are powerful tools that can help us understand and explore the dynamics of spatiotemporal behaviors, but only if they are an accurate match to reality. The goal in this paper is to use experimental data to build, validate, and test such a model. We are specifically interested in understanding how simple, local rules regarding food exchange, followed by all members of a bee colony, can lead to efficient, global food distribution. To that end, we began with a simple stochastic model of trophallaxis among self-propelled agents (i.e. individual bees) moving and interacting in a two-dimensional arena, constructed using NetLogo (Wilensky, 1999). A graphical representation of the model is shown in Fig. 1. Bee-agents can move freely in the simulation arena, which is a 33*×* 33 square box, with periodic boundaries.

The amount of food that agent *i* carries in the *n*^*th*^ time step is denoted as *f*_*i*_(*n*) and shown with gradations of red in the Figure. Agents are categorized into two different sets—*fed* and *deprived*—according to their initial food level: *f*_*i*_(0) = 1 and *f*_*i*_(0) = 0, respectively. At the beginning of each simulation, both groups are dispersed randomly across the simulation arena, with random initial headings. An agent can be in one of the two states at each time step during the simulation: *available*, which denotes that it is ready to initiate a food exchange interaction if it encounters another agent, or *busy*, which denotes that it is currently involved in a food exchange and cannot initiate another one. (For simplicity, we are only considering pairwise food exchanges in this model.)

We model the motion of each agent as a correlated random walk on the simulation arena. At each time step, every available agent modifies its previous heading by an angle increment Δ*θ*, drawn from a uniform distribution with mean *θ* = 0 and standard deviation *θ*^*^, and moves in that direction. That is, if *θ*^*^ = 0, agents can only walk in a straight path; if *θ*^*^ = *π*, agents may change their headings to any random direction at each time step. Algorithm 1 summarizes the action rules for each bee agent at each time step, beginning with each agent checking its immediate neighborhood (a circle of radius one) and performing the *θ*^*^-correlated random walk step to the targeted neighborhood that is described above. If that region is already occupied, the agent randomly changes its heading (“Redirect”) and continues its motion toward a non-occupied space in the neighborhood. If all neighboring space is occupied, the agent is not allowed to move; this effectively constrains the motion of the agents to 2D by disallowing superposition.

Trophallaxis exchanges in the model happen when available agents with different food levels encounter one another (“Find target”): if two available agents *i* and *j* are within one distance unit, and Δ*f* (*n*) =*| f*_*i*_(*n*) *-f_j_*(*n*) *|>* 0, then they initiate a food-exchange process. Both members of an encounter pair will then change their status to *busy* and stop moving until the exchange is complete. The exchange itself involves dividing the total food evenly between the two:

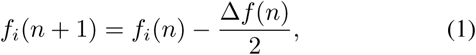

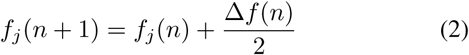

where Δ*f* (*n*) is the difference between the food levels of the two agents at the *n*^*th*^ time step.

The model runs until the food is distributed evenly across all agents, as measured with a two-part metric that assesses both the variance of the food among individual agents at each time step and the change in that variance over successive time steps. The variance is calculated in the following way:

### Algorithm 1

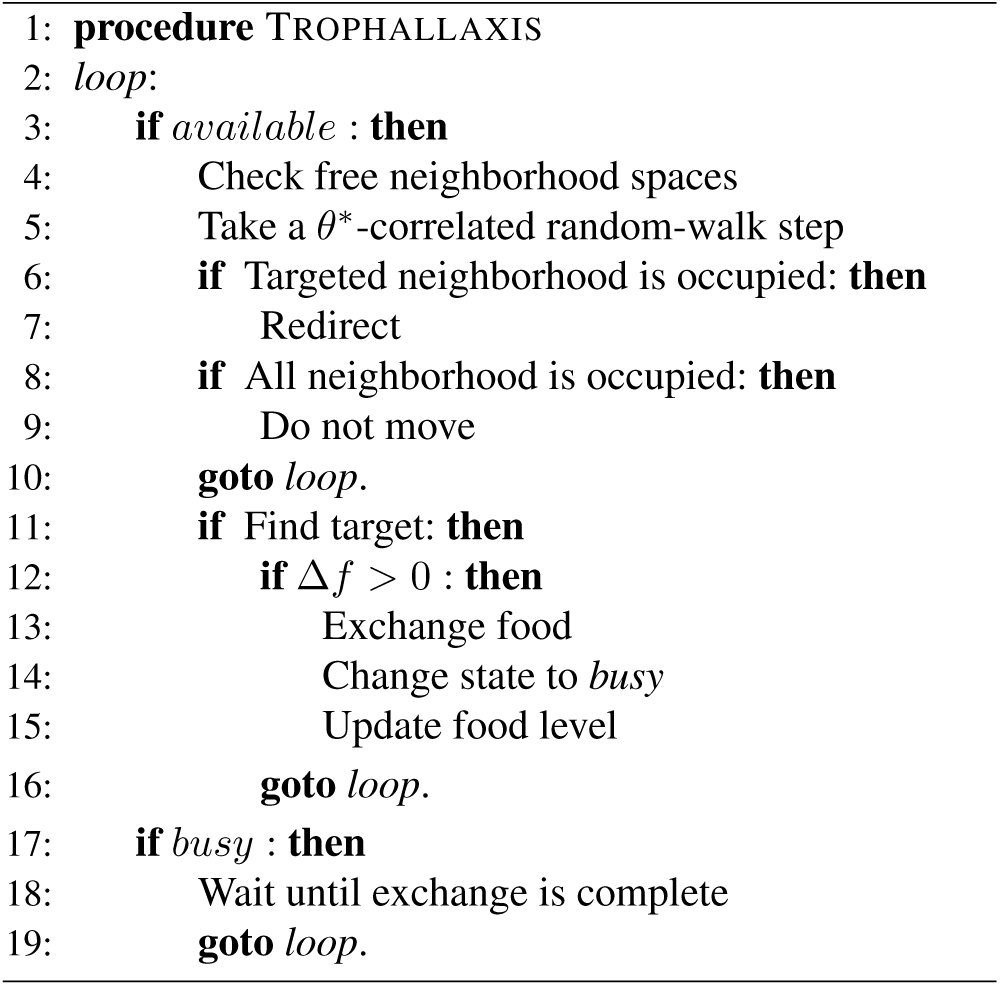

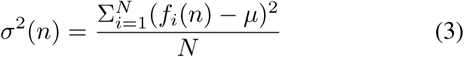

Where 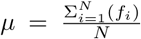is the average of the food across all agents in the system. The stopping condition is defined as when both of the following conditions are met:

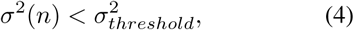

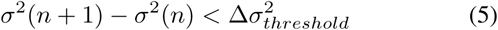

At the beginning of each model run, *σ*^2^(*n*) depends only on the fraction of fed bees. As the simulation progresses, the distribution of *f*_*i*_(*n*) values widens (i.e., *σ*^2^(*n*) increases) until reaching a plateau. The time to reach this plateau—the *convergence time*—is a measure of efficiency. To formalize its definition, we use the variance thresholds defined above, setting 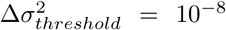 and 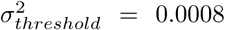. These choices ensure that there are no agents with *f* (*i*) = 0 at the end of the simulation run. Of course, the convergence time will depend on the values of the model parameters and the initial conditions (e.g., the density of bee-agents in the arena); we explore these effects in depth later in this paper.

## Laboratory Experiments

To finalize the design of the model and validate its results, we performed a set of laboratory experiments with honeybees, *Apis mellifera* L. Fig. 2 shows a top view of the experimental setup. The arena is a 33 *×*33*cm* square, covered with non-glare plexiglass to keep the bees contained and limit their motion to 2D. The subject bees were collected one day in advance and divided into two separate groups totalling about 120 individuals. One group was deprived of food for 24 hours before each experiment; the others had constant access to food. These fed bees, which comprised 5-10% of the whole population in each experiment, were carefully marked with a pink circle on their thorax. We started each experiment by introducing the deprived bees into the arena and allowing them 15 minutes of equilibration time before introducing the fed bees. Trophallaxis encounters began shortly thereafter. We collected videos with high temporal resolution of the arena using a camcorder (29.97 fps); for about 30 minutes in each experiment, beginning when the deprived bees were introduced into the arena.

The deprived bees were initially scattered around the arena, as shown in Fig. 2(a). After the fed bees were introduced, the bees began to move around and aggregate into clusters, as in Fig. 2(b). We also observed some interesting variations in the movement of the fed bees. To quantify these observations, we calculated values for a number of spatiotemporal metrics on the movement patterns, beginning with the tracks of the fed bees, which are easily computed using color thresholding techniques on the pink dots. The lengths of these tracks revealed that the fed bees moved quickly at first (Fig. 3(a)), then appeared to slow down (Fig. 3(b–c)), and then speed up again (Fig. 3(d–e)). These observations were used in validating some preliminary versions of the model.

**Figure 3:**
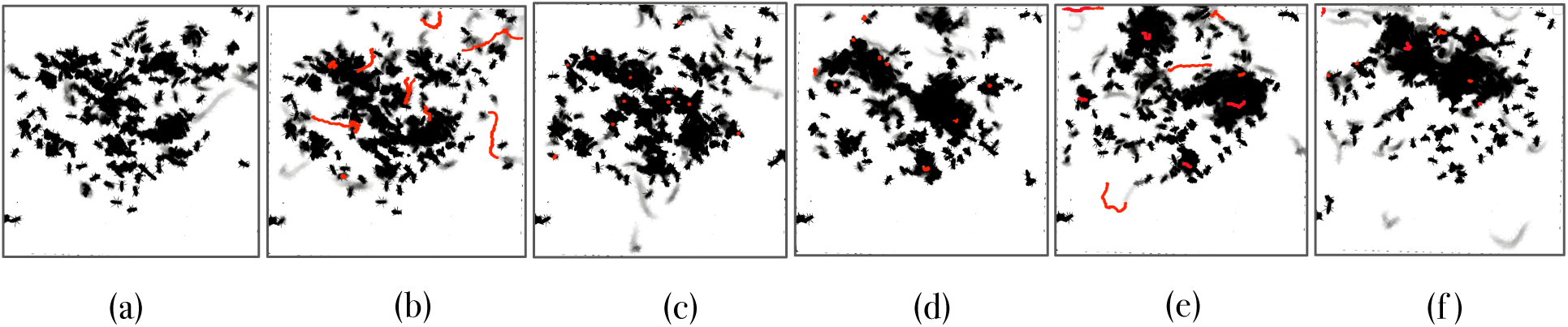
A sequence of snapshots of the average density maps, each constructed across 100 frames (i.e., 3.3 seconds), showing the temporal evolution of one experiment. The average density maps are useful proxies for the movement of deprived bees (the gray traces). The tracks of the fed bees during each period are shown in red. (a) Five minutes before the introduction of the fed bees. A temporal sequence: (b) 10 seconds, (c) one minute, (d) two minutes, (d) three minutes, (e) four minutes, (f) five minutes after the introduction of the fed bees.

Tracking the *unmarked* bees is a major challenge, however, despite recent improvements in tracking techniques (Boenisch et al., 2018; Bozek et al., 2018), because of the occlusions in the clusters, which makes it all but impossible to identify individuals. As a way around this problem, we converted the videos of each experiment to a sequence of *average density maps*, constructed using Otsu’s method (Otsu, 1979). This process thresholds the grey-scale image into black and white, separating the pixels into two classes: bee or non-bee (i.e. background). These plots—which are useful proxies for both the density and the motion of the unmarked bees in our experiments, as well as for the collective motion of the whole group—have been recently used for tracking people in crowd scenes with high density and heavy occlusions (Rodriguez et al., 2011; Wang et al., 2018).

We then used metrics from topological data analysis on these average density maps in order to quantify the cluster structure and track that structure over time. These calculations, which play a central role in our model validation procedure, are described in the following section. We also carried out a separate set of experiments with two pairs of deprived/fed bees in the experimental arena to measure the duration of trophallaxis encounters. This focused experiment allowed us to identify every individual food exchange encounter and get a precise measure of its duration. Most of these encounters were short; the longest was roughly 50 seconds. Fig. 4 shows a histogram of these durations.

**Figure 4:**
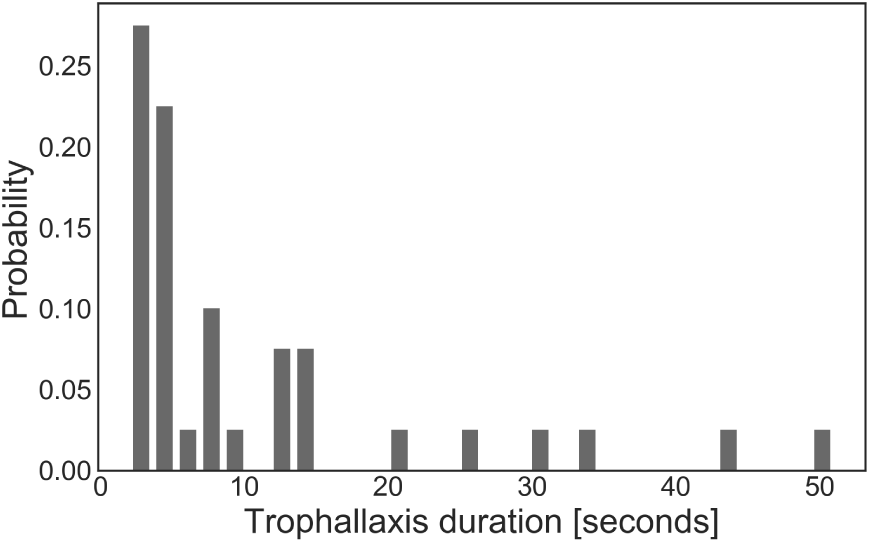
Histogram of trophallaxis duration in the experimental arena recorded over the course of an hour.

## A Data-Driven ABM for Trophallaxis

The experiments described in the previous section allow us to select physically realistic values for some of the important free parameters in our model, particularly the time required to exchange a given amount of food, which in turn defines the time scale in the simulations. It has been previously estimated that the time required for a trophallaxis exchange is proportional to the amount of food involved (Greenwald et al., 2015). We assume that fed bees are carrying as much food as they can, which we call 1.0 food units. That defines the maximal amount of food exchange in a single interaction in our model as 0.5 food units. We set the exchange duration for this amount of food to 50 seconds (i.e., 50 model time steps) to match the longest event in Fig. 4. We choose movement speed of the bee-agents to be one centimeter per second, consistent with experimental observations.

To validate this model, we focus our attention on the structures that form in the real and simulated groups of bees. There are of course many metrics for spatiotemporal analysis; based on our observations of the cluster structure in the experiments, we chose to use a method from computational topology called connected-component analysis. In general, the maximal connected subsets of a non-empty topological space are called the connected components of the space; every component is basically a closed subset of the original space. In computer-vision applications, a connected component refers to any group of spatially adjacent foreground (dark) pixels that are connected to each other via an eight-pixel measure of connectivity (i.e., all neighboring pixels in a 2D grid). We use the average density maps from our experiments to calculate the pixel area of a single bee and then use that value to identify the connected components^1^ (i.e., bee clusters) and estimate the number of bees in each one. We track the number and size of those clusters with time, and compare all of that information to corresponding values for the bee-agents in the simulations.

Fig. 5(a) shows a sequence of average density maps computed at one-minute intervals during one of our experiments, with the detected clusters on each image outlined in red. The dotted square in the middle figure in Fig. 5(a) corresponds to the minimum detected cluster size. Fig. 5(b) shows the trends in the number of clusters at each time step (*×*) and their average size (⊗) across the first six minutes of *all* experiments, with a vertical line indicating the time at which we introduced the fed bees into the arena. After this time, as Fig. 5(b) shows, the average number of bees in the clusters shows a clear increase. At the same time, the number of clusters reduces, suggesting that they merge as time passes. To quantify these relationships, we calculate the Pearson correlation coefficient between cluster sizes and counts to be −0.78. This strong negative cross-correlation confirms our observation that the bees do aggregate once the fed bees are inside the experimental arena—perhaps because this behavior supports the exchange of food.

**Figure 5:**
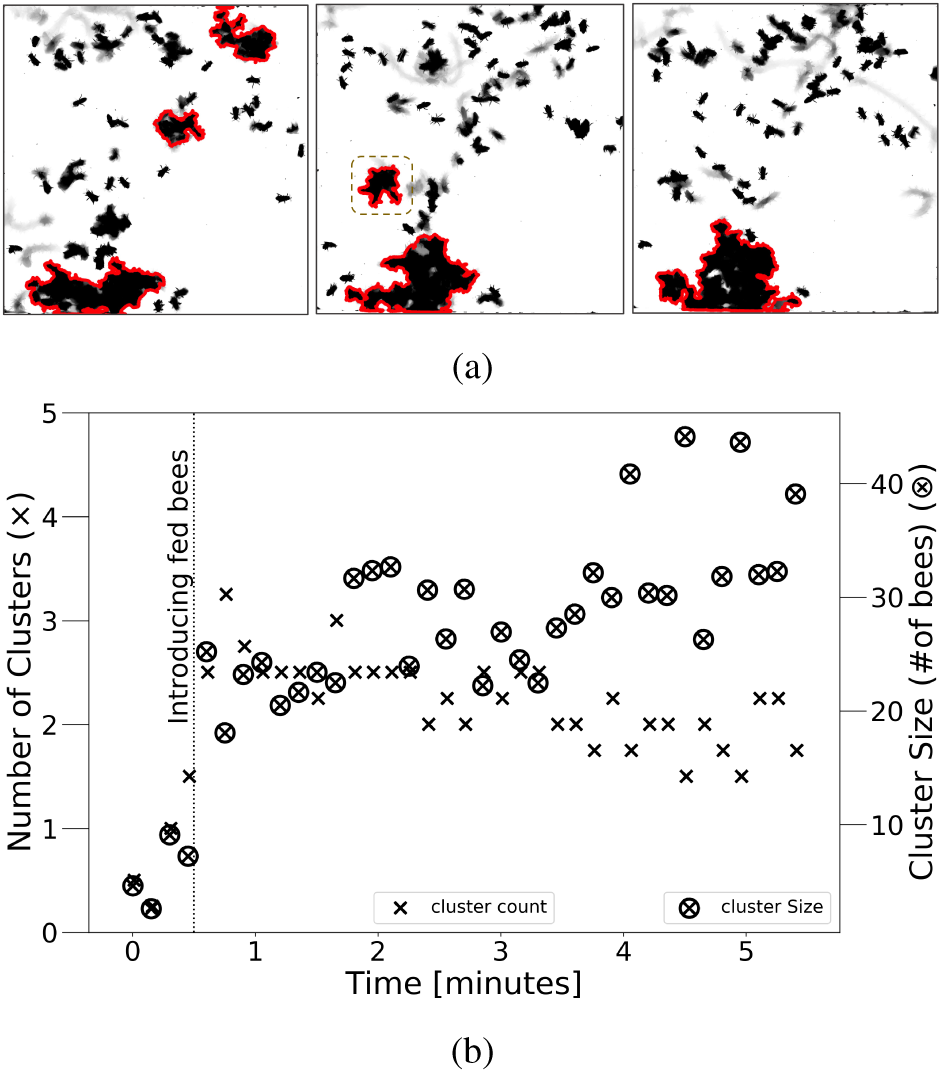
Cluster analysis of experimental data (a) Clusters detected in average density maps constructed at the first, second, and third minute of a representative experiment, shown outlined in red. The dotted square in the middle image highlights the smallest cluster that our algorithm searches for. (b) Temporal trends in the average size (⊗ signs) and the number of clusters (× signs) in all experiments after the full bees are introduced.

The bee agents in the first set of our simulations rarely form any kind of clusters. This is visually apparent from the model snapshot in Fig. 6(a) and also reflected in the detailed calculations shown in green in Fig. 7, which track the mean and variance in the cluster sizes over time. This is not surprising, since the model without attraction uses a simple random walk as a motion rule. Reasoning that the cluster formation observed in the experiments—the red trace in Fig. 7—may result from attraction between individuals, we modify the rules of motion in Algorithm 1 to include inter-agent attraction. Specifically, agents *i* and *j* that are within some *attraction range* (*r*) move toward one another one centimeter in the next time step:

**Figure 6:**
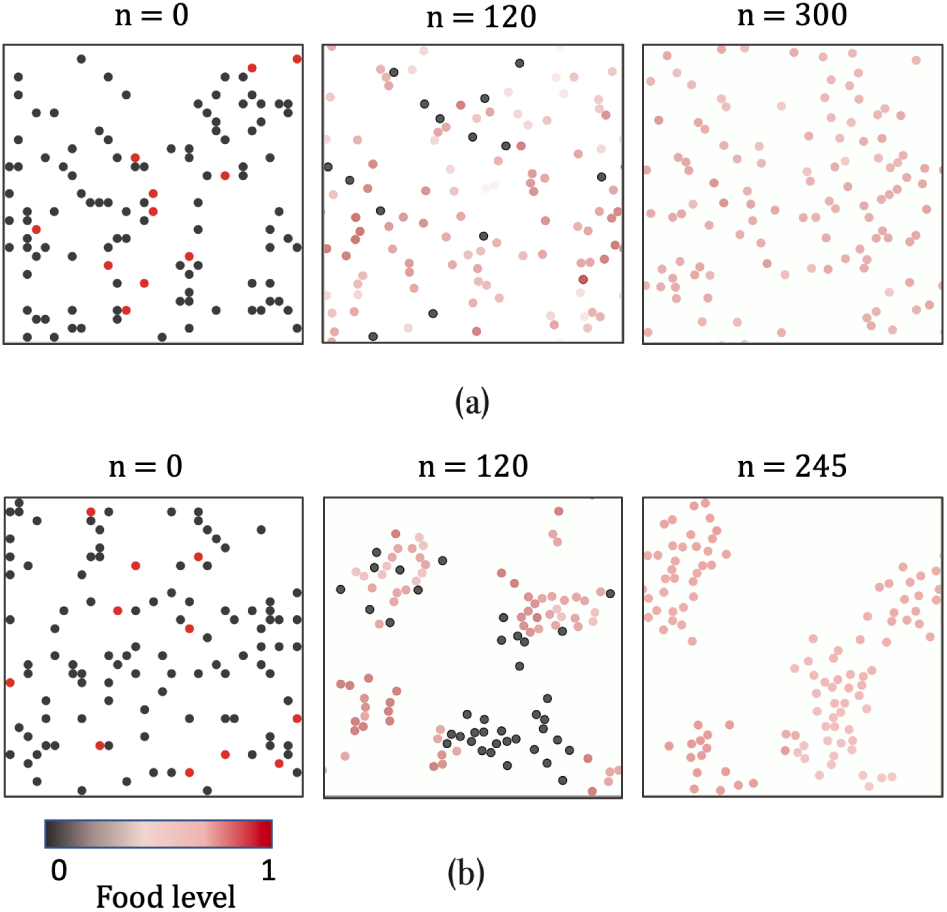
(a) Top row: snapshots of the agent-based model for trophallaxis without attraction, at three different time steps. (b) Bottom row: the same model with inter-agent attraction. As in Fig. 1, color reflects the food level of individual agents in both models. At the beginning of both simulations, the fed (dark red) and the deprived (gray) bees are scattered randomly in the simulation arena; the second image in each row shows an intermediate state of the model and the third image shows the state of the system, at the end of the simulation, when food is distributed evenly across all agents.

**Figure 7:**
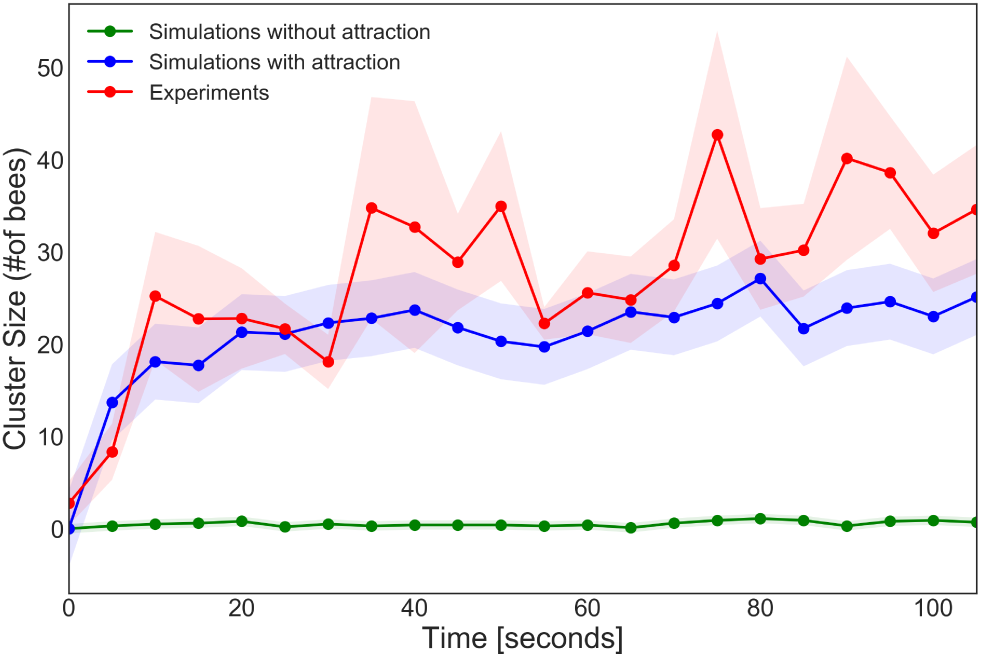
Comparison of the changes in cluster size in experiments and simulations with/without attraction. The shaded area represents the standard deviation from the mean.

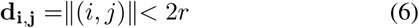

where, d_i,j_ is the Euclidean distance between the two agents. The value of attraction range can be in *r* = [0, 4], where *r* = 0 basically means there is no attraction involved and *r* = 4 assumes an attraction circle of radius four around each agent. As is visually apparent from the series of snapshots in the bottom row of Fig. 6, this modification caused clusters to form in the groups of bee-agents. This accordance between the spatiotemporal structure of the real and simulated bees is reflected in the better match between the red and blue traces in Fig. 7. To quantify this, we calculate the values of the normalized root mean squared error between the red (experimental) trace and the two model traces. For the model without attraction, this value is 0.87; for the model with attraction, it is 0.28. These values confirm that the model with attraction is a better match to the natural behavior of the bees in the experiments.

## Whence Efficiency?

Using this validated model, we then explored the effects of the attraction range (*r*) and the width of the turning angle of the random-walk (*θ*^*^) upon the efficiency of food distribution among the agents. Bee density is another important variable, as it can affect the frequency of interactions. Although not directly investigated in the experiments above, variations in bee density can guide future experimental designs. Bee density (*ρ*) can be operationalized as the number of bees divided by the area of the simulation box. Fig. 8 illustrates the change in convergence time as we vary this density between 0.01 and 0.1, averaged over 500 runs.

**Figure 8:**
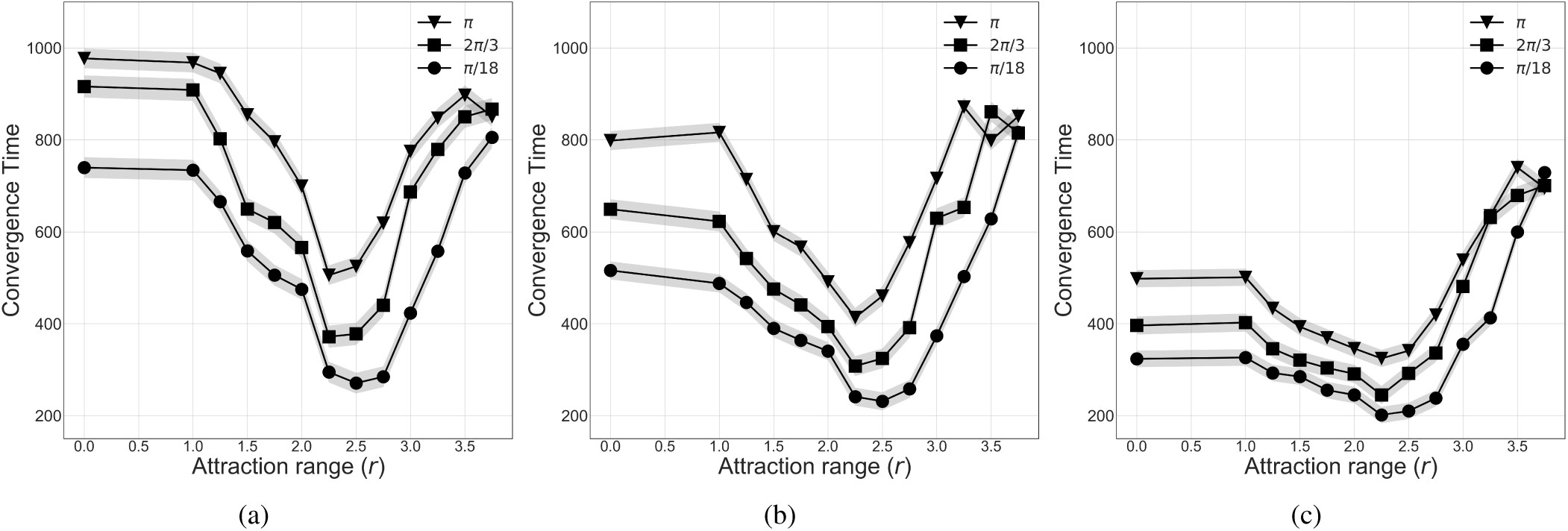
Simulation results for three different density and *θ*^*^ values as a function of attraction range (*r*). Convergence time for (a) small density (*ρ* = 0.01); (b) medium density (*ρ* = 0.05); (c) large density (*ρ* = 0.1). All results are averaged over 500 simulations and the shaded area represents the standard deviation from the mean.

As is clear from the three subplots in that figure, the average convergence time generally increases with *θ*^*^, regardless of the density. *That is, a more-tightly constrained random walk leads to more-efficient food distribution.* This has implications for understanding the local behavioral rules of individual fed bees. Specifically, a bee that has just finished feeding a nestmate faces a dilemma: who should it feed next? An intuitive solution would be to feed another nearby nestmate as quickly as possible. Our results suggest, however, that *not* feeding an immediate neighbor, but rather continuing her motion to reach a more-distant nestmate and share her food at a later time point, leads to a more efficient distribution of food. Recall that when *θ*^*^ = *π,* agents can change their heading in *any* direction at each time step. This encourages more local exchange events around the initial location of the donor agents. When *θ*^*^ is small, though, the model reveals that the probability of finding a local target is lower and that agents travel farther between trophallaxis interactions, leading to faster spatial distribution of food across the group.

Unsurprisingly, higher densities generally cause faster convergence, probably because of the associated increase in the encounter likelihood at each time step. Notice how the effect of the width of the turning angle on the convergence time dies out at higher densities and with larger attraction ranges. This is likely because higher attraction ranges lead to earlier formation of clusters—especially at higher densities, when agents are mostly within the attraction ranges of one another. Our simulation results also show that *short-range attractions, namely r* = (1, 3), *appear to increase the efficiency of food distribution among the agents, especially in lower densities.* Interestingly, the convergence time increases sharply when larger (i.e., non-local) attraction ranges are allowed, suggesting a less-efficient food distribution in these situations. This is probably due to early cluster formation, well before the food starts to spread. A potential side effect of this is the formation of clusters of deprived agents, whose members could block each others’ access to agents with food. This could lead to slower convergence of the model and less-efficient food propagation across the group.

Many of these assertions can be evaluated by calculating the number of encounters between the simulated bee-agents. Fig. 9 compares the average number of unique encounters in model runs with and without attraction. The higher number of unique encounters in the model with attraction increases the mixing between the two groups and leads to more-efficient food spread. (The results in Fig. 9 were calculated with *θ*^*^ = *π/*3 on a medium-density system, but the pattern in the encounter rates generalizes across all ranges of these parameters.)

**Figure 9:**
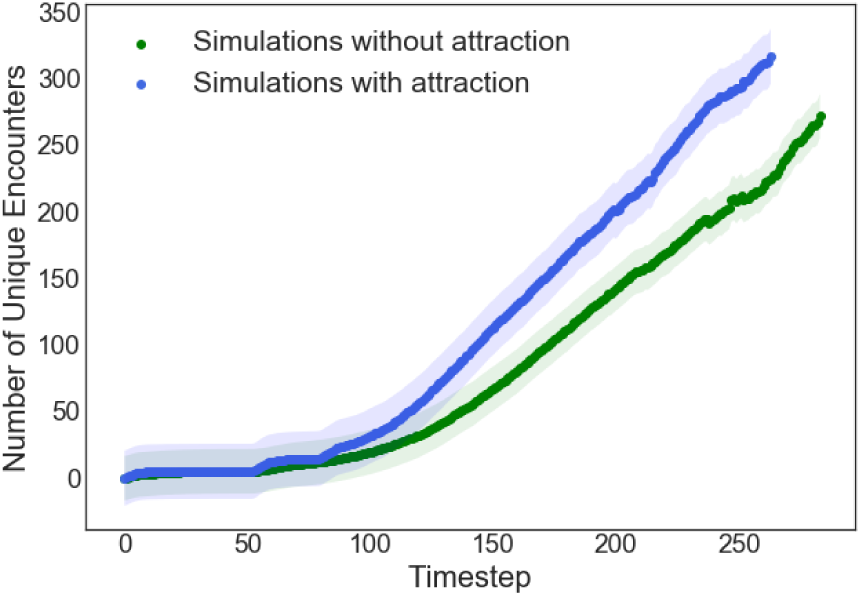
Average number of unique encounters at each time step during simulations with (*r* = 2, blue) and without (green) attraction.

## Discussion and Conclusion

Social insects and their collective intelligence have long been a reliable source of inspiration for the design of artificial multi-agent systems, optimization algorithms, and mostly swarm robotics (Bonabeau et al., 1999). Beyond its importance for modeling and understanding distributed feeding in biological systems, this study could inform engineering applications, such as electrical power sharing in swarms of search and rescue robots (Schioler, 2007). A well-known problem in autonomous swarm robotics, for instance, is re-charging of robots that are far away from a charging station (Schmickl and Crailsheim, 2006; Rubenstein et al., 2012). To prosper, a honeybee colony has to solve a similar problem; their solution involves forager bees collecting resources (food) and sharing it via trophallaxis, essentially charging nestmates who do not have direct access to those resources. The results of this study could potentially be used in designing more efficient, self-organizing systems using mutual power sharing, eliminating trips to the charging station and thereby increasing the efficiency of the overall system. As another example, the mutual charging system between electric cars can happen while they are on the move; effective strategies for charge-sharing among these vehicles could centralize and streamline the charging infrastructure for these important systems, which is both cost-efficient and leads to cleaner cities.

The agent-based model presented in this paper is not only inspired by trophallaxis behavior in some abstract way, but also designed and validated using laboratory experiments on honeybees. The rules in the model, and the values of the free parameters in those rules, were chosen via targeted experiments; the overall model was validated via comparisons between the movement patterns in the real and simulated bees. A comprehensive set of simulation results, performed with the validated model, suggest that the movement patterns of the individual agents, as well as their density and the range over which they are attracted to one another, affect the food-distribution efficiency. In particular, a combination of small turning angles, higher densities, and shorter-range attraction leads to the most efficient food distribution among the agents.

Our model makes testable predictions about optimality at different bee densities (*ρ*), and is currently guiding another set of experiments, with the long–term goal of studying the effects of short-range attractions on honeybees’ interactions. The focus of that work will be on low values of *ρ*, as our current model predicts a significant effect in this regime. We also plan to conduct more experiments with the focus on tracking the food and how it is being propagated among the bees. This would allow us to measure the food transfer rate via trophallaxis encounters, and make more robust and realistic connections between efficiency of food distribution in bees and convergence time in the model. This can be accomplished building on the technique used in (Greenwald et al., 2018) to study the trophallactic interaction networks in ants.

## Acknowledgements

This work was supported by the BioFrontiers Institute at the University of Colorado Boulder (internal funds). We would like to thank Charlotte Gorgemans, Gary Nave and members of the Peleg lab for insightful feedback and discussions.

## Footnotes

1 We choose to focus on the larger clusters: those containing at least 7-10 bees.

